# Optimal mating of *Pinus taeda* L. under different scenarios using differential evolution algorithm

**DOI:** 10.1101/2022.07.18.500513

**Authors:** Khushi Goda, Fikret Isik

**Affiliations:** Genetics Program, North Carolina State University, Raleigh, NC, USA; North Carolina State University Cooperative Tree Improvement Program, Raleigh, NC, US

**Author notes:** **Corresponding author:** Fikret Isik. **Contributions** Goda developed the software, planned the study, analyzed the data, and wrote the first draft of the manuscript. Isik conceptualized the study and contributed to the writing of the manuscript. Both authors read and approved the final version of the manuscript. **Competing interests** The authors declare no competing interests. **Funding** North Carolina State University Cooperative Tree Improvement Program and USDA-NIFA – Genomic Selection in Forest Trees: Beyond Proof of Concept (PD: Isik) (award # 2019-67013-29169).

**Keywords:** optimal mating, monoecious species, optimization, inbreeding, coancestry, differential evolution

## Abstract

A newly developed software, AgMate, was used to perform optimized mating for monoecious *Pinus taeda* L. breeding. Using a computational optimization procedure called differential evolution (DE), AgMate was applied under different breeding population sizes scenarios (50, 100, 150, 200, 250) and candidate contribution scenarios (max use of each candidate was set to 1 or 8), to assess its efficiency in maximizing the genetic gain while controlling inbreeding. Real pedigree data set from North Carolina State University Tree Improvement Co-op with 962 *Pinus taeda* were used to optimize objective functions accounting for coancestry of parents and expected genetic gain and inbreeding of the future progeny. AgMate results were compared with those from another widely used mating software called MateSel (Kinghorn, 1999). For the proposed mating list for 200 progenies, AgMate resulted in an 83.7% increase in genetic gain compared with the candidate population. There was evidence that AgMate performed similarly to MateSel in managing coancestry and expected genetic gain, but MateSel was superior in avoiding inbreeding in proposed mate pairs. The developed algorithm was computationally efficient in maximizing the objective functions and flexible for practical application in monoecious diploid conifer breeding.

**Study implications:** A dataset from a breeding population of loblolly pine (*Pinus taeda* L.) was analyzed using an optimal mating software, AgMate (developed by the authors), to optimize the selection, contribution, and mating of candidates simultaneously. The software helps breeders make decisions on which tree to cross with which tree and how many times, such that the trees are not related to each other and would result in the best performing progenies. AgMate is effective in meeting the breeding objectives for monoecious species. The open-source, easy-to-use, and flexible AgMate software, also available as a website, is invaluable in helping breeders to create optimal matings for future generations, which balance the pursuit of maximizing genetic gain while maintaining genetic diversity.

## Introduction

Conifer tree breeding programs face the ever-present challenging goals of managing genetic diversity while breeding for genetic gain (Isik and McKeand 2019). This problem is exacerbated for conifers, like *Pinus taeda*, which suffer from inbreeding depression (Ford et al. 2015; Franklin 1972; Hedrick, Savolainen, and Kärkkäinen 1999; Remington and O’Malley 2000). Conifers are highly fecund species, and the problem arises when high selection intensities are used, and a limited number of best-performing individuals are responsible for genetic contributions to the next generation (Hamilton, 2020). This leads to a rapid increase in the inbreeding levels in the finite breeding population (Isik and McKeand 2019). Numerous studies have reported the effects of inbreeding on conifers, such as reduced fecundity, decreased growth, and low survival rate (Durel, Bertin, and Kremer 1996; Williams and Savolainen 1996). The reduction in genetic value due to inbreeding is called inbreeding depression (Falconer and Mackay 1996; Lynch and Walsh 1998), and this has detrimental effects on the long-term response to selection.

Selection and mate allocation of candidates where the genetic gain is maximized while incorporating some form of management for inbreeding in the long-term breeding population can be shaped as an optimization problem. Several methods have been proposed to solve this optimization problem. Meuwissen’s (1997) Optimum Contribution (OC) Selection aims to find maximize genetic gain by optimizing the candidate contribution under a constrained rate of inbreeding (Hallander and Waldmann 2009; Kerr, Goddard, and Jarvis 1998; J. Woolliams et al. 2015; John Woolliams and Thomson 1994). Various methods have been applied to solve the OC problem: Lagrange-multipliers (Dagnachew and Meuwissen 2016; Meuwissen and Sonesson 1998), semi-definite programming (Kerr et al. 2015; Pong-Wong and Woolliams 2007) and second-order cone programming (Yamashita, Mullin, and Safarina 2018). While the OC provides a solution to the contribution optimization problem, it does not offer a solution to the mate allocation problem. An evolutionary algorithm, like the differential evolution algorithm (DE) by Storn and Price (1997), has been shown to optimize for candidate selection and contribution simultaneously, solving the mate allocation problem as well (Kinghorn & Shepherd, 1999). DE effectively optimizes multiple goals simultaneously, a property often absent in other algorithms (Price, Storn, and Lampinen 2006). Numerous studies have shown the effectiveness of adopting DE into their breeding programs for various species (Carvalheiro, Queiroz, and Kinghorn 2010; Isik and McKeand 2019; Kerr et al. 2015; Weigel and Lin 2000; Yoshida et al. 2017). The application of DE to breeding programs was initially developed for animals, especially livestock breeding, and remains not well-suited for many forestry species (Hamilton 2020). Modifications to preselection and mate selection algorithms successfully helped manage the increase in average coancestry over time when planning future mating designs for overlapping generations in the forest trees (Kerr et al. 2015). Isik and McKeand (2019) showed the effectiveness of using DE designed for animal breeding in planning the 4th cycle of *Pinus taeda* breeding. A moderate balance between genetic gain and parental coancestry scenario suggested that 63% of the genetic gain could be obtained with a slight increase in the average inbreeding level. To our knowledge, no software has been specifically designed to use DE in monoecious *Pinus taeda* breeding programs. To this point, we developed new software, called AgMate: An Optimal Mating Software for Monoecious Species (Goda and Isik 2022), to bridge the gap.

The study analyzes different scenarios using an empirical data set to evaluate the ability of AgMate to meet the optimizing objectives. The study also aims to show a proof-of-concept of the effectiveness of AgMate by comparing the results from different mating scenarios with MateSel software by Kinghorn (1998), a widely used software by North Carolina State University Tree Improvement Program.

## Materials and Methods

### Empirical data set

North Carolina State University Cooperative Tree Improvement Program (TIP) provided the empirical *Pinus taeda* data set used in this study. TIP has managed a pine breeding program for over 66 years and has a comprehensive database of the *Pinus taeda* breeding population. For this study, we used the Atlantic Coastal Plains Breeding population of 962 trees selected to establish the fourth cycle population. Volume-Straightness-Rust (VSR) index breeding value (IBV) was calculated for each candidate by putting proportional weights on each trait of interest. The traits of interest mainly concerned wood properties, namely volume of wood, straightness of the wood, and rust resistance ability of the wood. IBV = 0.6*Volume + 0.2*Straightness -0.2*Rust, where the weight on desirable traits of volume and straightness are positive 0.6 and 0.2 respectively, however negative 0.2 for undesirable rust trait. The negative weight on rust ensures that candidates with higher breeding values for rust, that is, genotypes with a higher probability for rust disease, are penalized. Individuals with IBV>1 were selected as candidates for the mating designs (Isik and McKeand 2019). The 962 candidates formed the input candidate list for AgMate to design mating lists of 50, 100, 150, 200, and 250 mate pairs. The input file for AgMate is a CSV file with five fields: CandidateID, Parent 1, Parent 2, Index, and MaxUse. Two additional scenarios of different MaxUse for each candidate were analyzed, AgMate_U1 for MaxUse set at 1 and AgMate_U8 for MaxUse set at 8.

### AgMate: An Optimal Mating Software for Monoecious Species

AgMate (Goda and Isik 2022) is a multifunctionality automated software developed to optimize the selection and contribution of candidates to create optimal mating lists. The overview of the software is outlined in Figure 1. AgMate is developed in R, and the detailed R code is available at https://github.com/khushigoda/AgMate. The ShinyApp version of the software is available here: https://khushigoda.shinyapps.io/AgMate/. The software handles the entire optimization process of completing the candidate list, calculating the relatedness between candidates, candidate selection and contribution decision, and eventually creating an optimized mating list. We discuss the processes present as functionalities in the software in depth below.

**Figure 1 (Goda and Isik, 2022).**
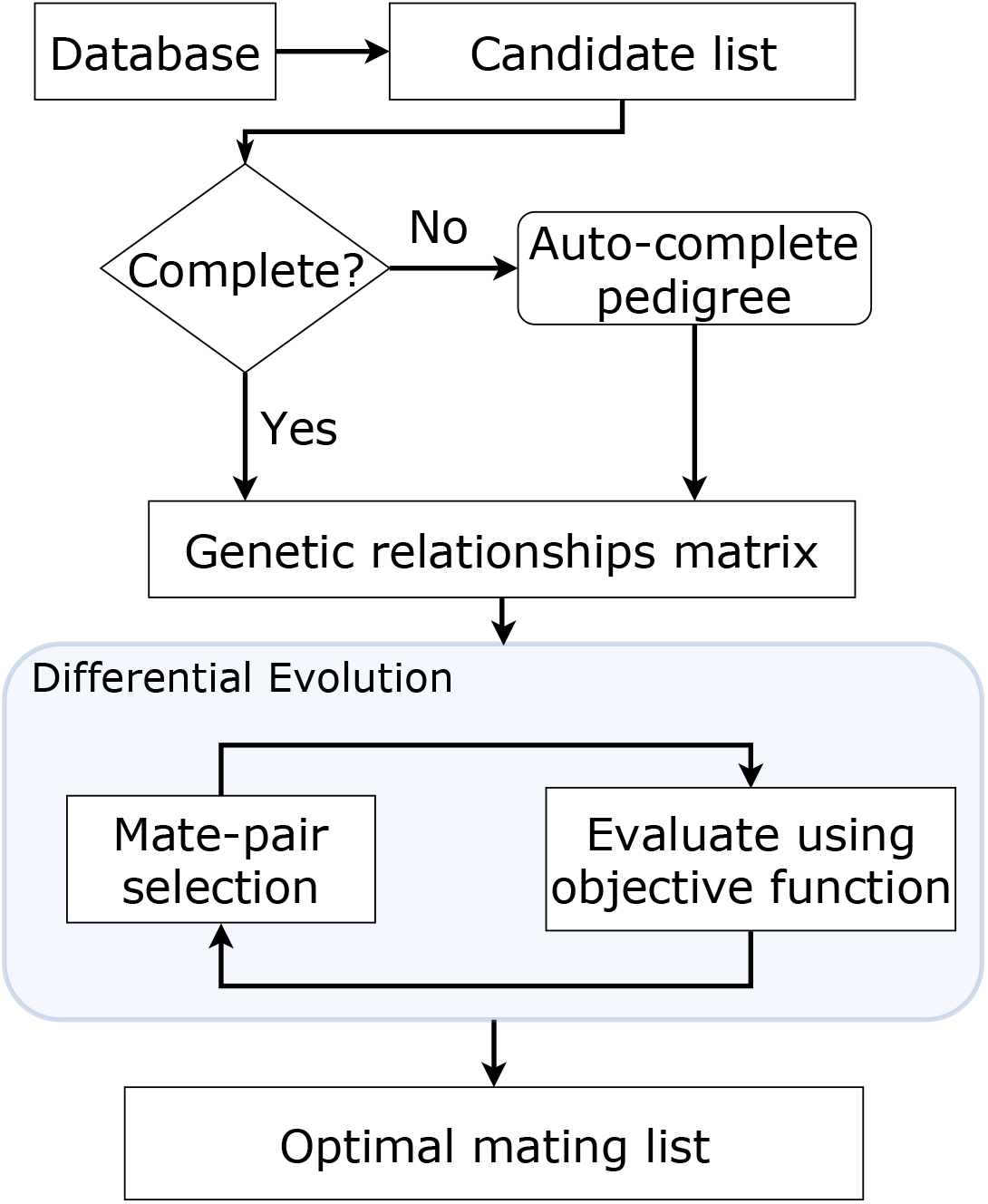
Schematic outline of AgMate: an optimal mating software for monoecious species. TIPRoot Database serves as the reference. The pedigree is automatically completed and ordered for preparing the candidate list input file. The numerator relationship matrix is calculated for the complete pedigree. Using a differential evolution algorithm, all possible mate pairs are searched and compared to produce numerous mating lists. These mating lists are evaluated based on an objective function that considers the total expected genetic gain from progeny, the mean inbreeding levels in progeny, and the coancestry between the selected parents. The best ranking mating list which satisfies all constraints is chosen as the optimal mating list.

### Auto-completion of pedigree

The first step of AgMate software involves the auto-completion of the pedigree. Two lists of individuals are required for this step: a candidate list, i.e., the list of individuals to be mated (962 candidates for this study), and a reference list, i.e., a list of all known individuals in the pedigree and beyond. The auto-completion process works by referencing the candidate list to the reference list using a quick search and sort method. This functionality aims to look up all known ancestors of the candidates from the reference list and add them to the candidate list, such that the parents always precede the progeny to complete the pedigree. North Carolina State University Cooperative Tree Improvement Program database called TIPRoot (Anonymous 2016) was referenced to autocomplete the pedigree of the 962-candidate list. Without an ordered and completed pedigree, essential relatedness information would not be considered in the downstream analysis of optimization of mate pairs accounting for inbreeding and coancestry. This is a crucial step to remove the redundant work of manually populating the candidate pedigree. The candidate file for AgMate is a CSV file with four columns: Candidate ID, Parent 1, Parent 2, EBV, and MinUse. MinUse for each candidate was set at 1. The reference file for AgMate is a CSV file with three columns: Individual ID, Parent 1, and Parent 2. Note that Candidate ID and Individual ID need to match for this process.

### Auto-calculation of numerator relationship matrix

Relatedness parameters between candidates are calculated using the numerator relationship matrix (**A** matrix). **A** matrix provides information about the coancestry, *θ* (Wright 1922) between any two candidates and the inbreeding, *F*_*x*_ (Falconer and Mackay 1996) level of each candidate. The completed and ordered pedigree of the 962-candidate list was used to compute the **A** matrix using Henderson’s recursive method (Henderson 1976; Mrode 2014) via AGHmatrix R package (Amadeu et al. 2016). The package functions to calculate the genetic relationships are part of AgMate software. A stringent coancestry constraint of 0.075 (*a*_*ij*_ = 2*θ* = 0.15) was set to prevent mate pairing between related candidates and avoid a rapid accumulation of coancestry in the mating population. This constraint does not allow any full-sibs, half-sibs, or more highly related candidates to mate with each other. Additionally, it also prevents self-crossing and reciprocal crosses, ensuring long-term management of inbreeding due to the severe effects of inbreeding depression on *Pinus taeda* (Ford et al. 2015).

### Optimization of matings in AgMate software: Differential Evolution

The differential evolution (DE) algorithm, an evolutionary algorithm (Storn and Price 1997), was applied to create an optimal mating list. The DE was modified and adopted by Kinghorn to optimize matings in animal breeding. Here we modified the methods proposed by Kinghorn to adapt the optimization methods for monoecious species like *Pinus taeda* and incorporate them into AgMate. DEoptim (Mullen et al. 2011), an R package, was incorporated into the AgMate software to run the DE analysis. The main modification to the optimization method was redesigning the mating matrix, a required setup before the optimization begins. Kinghorn initially proposed a mating matrix that was more adapted to species with male and female sexes (Brian P Kinghorn 2011), which is unsuitable for monoecious *Pinus taeda*. While the applications and framework of the differential evolution algorithm are outlined in the literature (Carvalheiro et al., 2010; Isik & McKeand, 2019; Kinghorn, 2011; Kinghorn & Shepherd, 1999; Yoshida et al., 2017), here we outline the processes incorporated into AgMate for the DE optimization of mating lists.

Candidate selection and the number of contributions per candidate are decided simultaneously by DE in AgMate software. According to the number of contributions of individual candidates, the candidate list is expanded such that there are duplications of candidates; each contribution gets a specific mention. For example, if a candidate has three contributions, the candidate will now be present thrice in the list, with each candidate being used only once (with repetition). This ensures that only the candidates to be mated are represented in the mating matrix and that each candidate with minimum use of more than or equal to one is mated. To initiate the mate allocation process, each candidate in the expanded candidate list is assigned a ranking criterion (Kinghorn, 2011), i.e., a random positive integer. The list is ranked based on this ranking criterion. This number has no link to the candidate’s breeding value or fitness. A mating matrix is designed with the expanded candidate list, where the rows are candidates ranked based on the ranking criterion and the columns are non-ranked candidates. Since the MinUse for each candidate was set at 1 for AgMate analysis, no duplicate candidates existed in the mating matrix.

Decisions regarding allowed mate pairing are based on the relatedness observed between the candidates. An initial mating list is created by allocating the best-ranked candidate, based on the ranking criterion, to the first available candidate that it has permission to mate with, and this process continues down the list till all the required mate pairs are created (explained elsewhere in (Gondro and Kinghorn 2008; Kerr et al. 2015). The DE optimization process then works iteratively to search the entire search space to create numerous such mating lists before finding the optimal mating list. Additional checks ensure that no two mate pairs are repeated (i.e., no full-sibs progeny) and the total number of mate pairs is adhered to. For this study, since the proposed progeny were limited to 50, 100, 150, 200, and 250 only, no candidate was allowed to be used more than once (i.e., no half-sibs progeny). The robustness of the mating lists is evaluated based on an objective function, as suggested by Kinghorn (1999). DE aims to find an optimal solution to the objective function (Eq.1), such that the mating list proposed would maximize genetic gain while minimizing the increase in progeny inbreeding and the coancestry between suggested mate pairs.

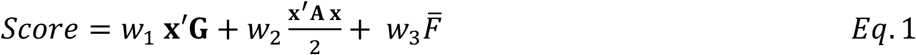

where ***x*** is the contribution vector of the parents, ***G*** is a vector of the parent’s estimated breeding values (EBVs), **A** is the numerator relationship matrix derived from pedigree, and 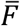 is the mean inbreeding coefficient in progeny. **x**^′^**G** gives the total genetic gain expected from the progeny, 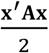 is the coancestry of selected parents; and *w*_1_, *w*_2_ and *w*_3_ are 1, -10 and -1 weights applied to progeny genetic gain, coancestry of selected candidates and progeny inbreeding, respectively. The negative weights result in mating lists with high progeny inbreeding levels and parental coancestry having a low score value. Thus, through the recursive DE algorithm, after searching the entire search space for possible mating lists and ranking them based on their score value, only the highest-ranking mating list which satisfies all constraints will be selected as the optimal mating list. AgMate scenarios were terminated after 10 DEoptim iterations. The final mating list output gives the proposed mate pairs, the expected mid-parent breeding values of their progeny, and expected inbreeding levels for each suggested progeny.

### Optimization of matings in MateSel software

Here we validate the AgMate software with the licensed version of MateSel (Kinghorn, 2011) (http://matesel.une.edu.au/) used at the North Carolina State University Tree Improvement Program. MateSel applies the same differential evolution (DE), an evolutionary algorithm developed by Storn and Price, 1995, to create the optimal mating list. For the MateSel software, the input file is a CSV file with five fields: CandidateID, SireID, DamID, Index, Maxuse, and Sex. MateSel requires an input of the completed pedigree of candidates, thus, the completed candidate list created by AgMate was used. MateSel re-orders the pedigree to ensure that the parents always precede the progeny. **A** matrix calculation takes up much of the runtime due to the sheer size of the matrix, especially when deep pedigrees are involved and the completed pedigrees are large. MateSel uses Jacques Colleau’s (Colleau 2002) faster method of calculating **x**^′^**Ax**, which is the mean parental coancestry, compared to simple matrix manipulations on the non-zero elements of the **A** matrix. The coancestry cut-off for MateSel analysis was set at 0.075, keeping it consistent with AgMate parameters. Since MateSel doesn’t allow MaxUse for bisexual candidates to be set at 1, the only scenario analyzed was MaxUse at 8. MateSel aims to optimize the objective function explained above as well but uses a frontier curve to come to an optimized conclusion. Weights for the objective function were kept constant at 1, -10, and -1 for *w*_1_, *w*_2_, and *w*_3_ respectively. MateSel was run under different scenarios to output optimal mating lists for 50, 100, 150, 200, and 250 mate pairs. For termination, MateSel was allowed to converge to 99.9% and then manually stopped. Optimization by DE in AgMate adopts this method with necessary enhancements for monoecious diploid *Pinus taeda* breeding.

## Results

The AgMate software successfully completed and ordered the pedigree of candidates by referencing the database and calculating the necessary distribution statistics. The completed pedigree had 1073 individuals for the initial population of 962 candidates. The average EBV, average coancestry coefficient, and average inbreeding coefficient of the completed pedigree of the candidate list were 3.9, 0.0015, and 0.0003, respectively. A check of the completed pedigree by referencing the NCSUTIP TIProot database showed no incorrect or missing information. AgMate was also used to complete the proposed optimal mating lists full pedigree for all scenarios, including those by MateSel.

AgMate_U1, where the MaxUse for each candidate is set at 1, was able to select more candidates to contribute to the next generation compared to AgMate_U8 and MateSel while resulting in a similar increase in the average index and coancestry level as MateSel (Figure 2 and 3). AgMate_U8, where the MaxUse is set at 8 for each candidate, used the least number of candidates to produce the required mate pairs (Figure 5). For the mating list of 50 mate pairs, the complete pedigree consisted of 206 individuals for AgMate_U1, 72 individuals for AgMate_U8, and 148 individuals for MateSel. By allowing a higher number of use cases for each candidate, AgMate tends to allow fewer candidates to create more mate pairs if the constraints on the coancestry of selected parents are not broken, and the objective function is still optimized.

**Figure 2.**
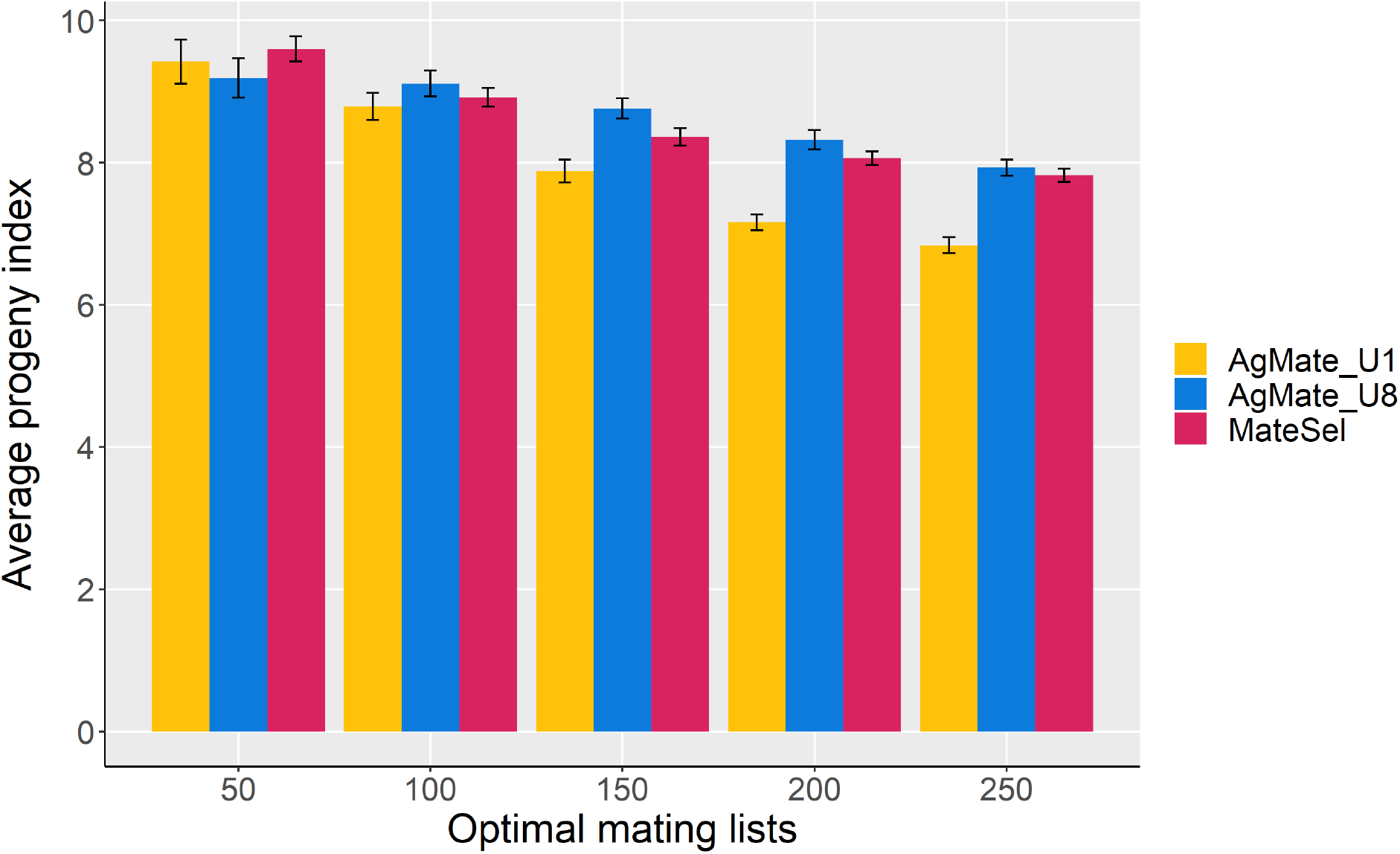
Comparison of AgMate and MateSel for progeny index under different mating lists (50 to 250). Three scenarios: AgMate with MaxUse set at 1 (AgMate_U1), at 8 (AgMate_U8) and MateSel with MaxUse set at 8 are shown. MateSel does not allow MaxUse to be set at 1 under bisexual conditions. The candidate population had an average index of 3.9, and all the mating lists scenarios showed improvement and resulted in a similar average progeny index.

**Figure 3.**
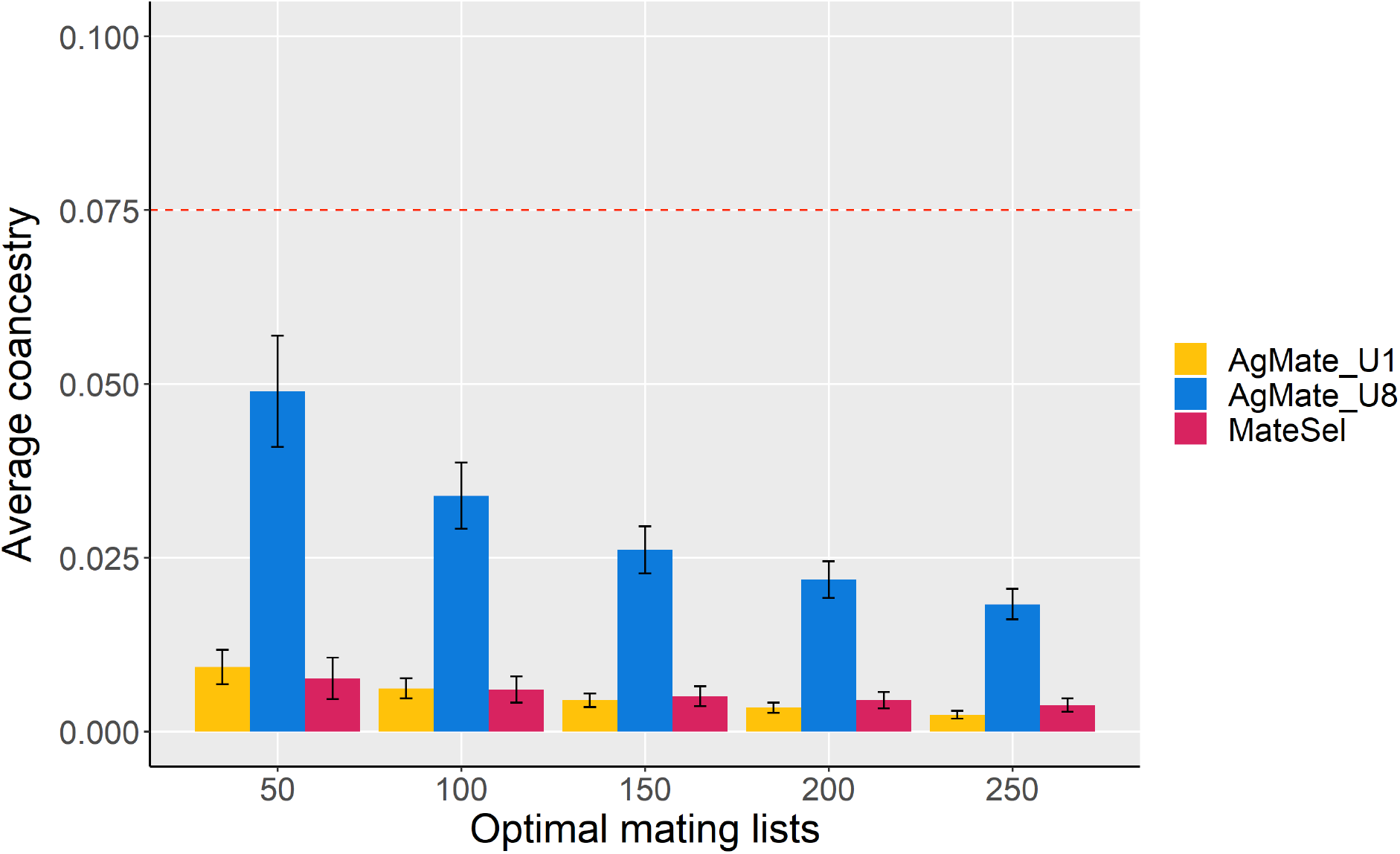
Comparison of AgMate and MateSel for average coancestry under different mating lists (50 to 250). The average coancestry is calculated for complete pedigrees. Three scenarios are shown: AgMate with MaxUse set at 1 (AgMate_U1), 8 (AgMate_U8), and MateSel with MaxUse set at 8. AgMate_U8 produced the highest average coancestry. AgMate_U1 and MateSel resulted in a minor increase in average coancestry and were comparable. No scenario broke the coancestry constraint of 0.075 (red line).

Reducing the number of allowed MaxUse of candidates allows AgMate to create more genetically diverse optimized mating lists, which are also optimized to minimize the increase in inbreeding and coancestry levels.

AgMate_U1 performs better than AgMate_U8 in the management of coancestry (Figure 3). As multiple candidates are used more than once in AgMate_U8, the overall relatedness in the optimized mating list increases because of an increased number of half-sibs in the proposed mate pairs. While the selected parents in the mating lists still follow the constraints and have coancestry lesser than 0.075, the average coancestry levels in the complete pedigrees are bound to increase every time a candidate contributes as a parent to more than one progeny. AgMate_U1 performed similarly at maintaining the increase in the average coancestry as MateSel under the same constraints and penalty on coancestry. Both software resulted in an increase in the average rate of coancestry, calculated for the entire pedigree of the proposed mating list consisting of the selected parents to be crossed, their progenies, and their ancestors over the different scenarios (Figure 3). For AgMate_U1, the increase in average coancestry when designing a mating list of 250 mate-pairs was 0.00242, and the increase for the same scenario with MateSel was 0.00381, which was only 36.5% more than AgMate_U1. For other scenarios of proposed mate pairs, MateSel and AgMate_U1 performed similarly (Figure 3). On the contrary, AgMate_U8 resulted in approximately 80% higher levels of coancestry than MateSel for the same MaxUse case of 8 for each candidate, but the average coancestry levels were still within the constraints (Figure 3).

All three scenarios, AgMate_U1, AgMate_U8, and MateSel, resulted in a similar average genetic gain, as measured by average mid-parent values of proposed mate pairs, with minimal differences. Overall, AgMate_U1 provided the least genetic gain of all scenarios, and AgMate_U8 had a high average progeny index, similar to MateSel. The mating list for 100 progenies designed by AgMate_U8 had the highest genetic gain of 57%, compared to the candidate list, while AgMate_U1 and MateSel had about 56%. While AgMate_U8 resulted in a slightly higher genetic gain for the proposed mating lists, the substantially higher increase in coancestry in these mating lists is unfavorable. AgMate_U1 provided a more balanced increase in genetic gain and management of increase in coancestry.

For the same level of genetic gain, AgMate_U1 created more diverse mating lists than AgMate_U8 and MateSel. On average, AgMate_U1 had 36% more individuals in completed pedigrees for the mating lists than MateSel. For example, on the proposed mating list of 150 progenies, the completed pedigree, which includes parents, ancestors, and progenies, was 575 for AgMate_U1, whereas it was only 363 for MateSel. The varied mating lists differed not only on the selected candidates but also in the contributions of the commonly selected candidates. For example, to generate the 150 mate pairs from the 300 unique candidates selected by AgMate_U1, only 99 were in common with MateSel. MateSel uses only 115 unique candidates to create a mating list of 300 candidates for 150 mate pairs. Out of these 115 candidates, 75 are used more than once, with one of the candidates being used a total of eight times (Figure 5). Candidates used multiple times tend to have higher EBVs, thus providing more genetic gain to the resulting mating lists. However, they also had an increased population of half-sibs in the mating lists, resulting in substantially higher coancestry in the mating population.

MateSel is highly efficient in preventing inbreeding in the progenies. MateSel reported zero inbreeding levels in the progeny for most of the proposed mating lists. AgMate, on the contrary, had higher inbreeding levels for all the scenarios. Since AgMate aims to use more unique candidates, it is bound to use candidates with a certain level of inbreeding. Figure 2 shows the number of mate pairs contributing to the increased inbreeding levels. For a mating list of 50 progenies, only 1 mate pair had inbreeding and was contributing to the average inbreeding level. A more stringent negative weight on the inbreeding level would help reduce the average inbreeding in AgMate.

## Discussion

Two different software, AgMate (Goda and Isik 2022) and MateSel (Kinghorn, 2011), both utilizing the DE algorithm (Storn and Price 1997), were applied for the optimization of candidate selection and contribution to design mate-pairs and create optimal mating lists for breeding of *Pinus taeda*. AgMate was applied under different scenarios in real data set from a *Pinus taeda* breeding population and validated by a routinely used and powerful software MateSel. Both the AgMate and MateSel software showed to be computationally efficient. The computational efficiency came mainly from using sparse matrices and optimized approaches to computing coancestry from numerator relationship matrices. AgMate utilizes Henderson’s recursive method (Henderson 1976; Mrode 2014) to calculate the numerator relationship matrix and to calculate coancestry from the completed and ordered pedigree of the candidates. MateSel uses Colleau’s (2002) indirect approach to compute coancestry. Both these methods help in the reduction of analysis run-time substantially. For example, the analyses by AgMate of the *Pinus taeda* data set, which had 1073 individuals in their full pedigree, which would have a numerator relationship matrix of size 1073 × 1073, ran in approximately 5 to 10 minutes (depending on the size of the mating list to create) on a PC with an Intel® Core™ i7 2.3GHz processor and 16GB RAM. Run-times by MateSel were equivalent to AgMate for the scenarios in this study, and since the manual stop option was exercised, run-times were not recorded.

While AgMate is computationally efficient, it also allows breeders the flexibility to optimize their mating lists on the factors they deem of importance by deciding on the weights to place on different components of the objective function. AgMate adopts the same objective function as MateSel (Kinghorn et al., 1999) and requires the breeders to input the weights to be applied for each component of the objective function. The flexibility of setting the weights of the objective function components allows the users to determine the ideal weights such that the optimization is based on the needs of the breeding program. Breeders need to consider the size of the population, the level of inbreeding and coancestry present in the breeding population, and the desired level of genetic gain, balanced with the desired level of inbreeding and coancestry in the future populations, with consideration for the management of genetic diversity. Knowing their breeding populations, users are more informed and better able to find the ideal weights that strike the appropriate balance between genetic gain and the coancestry of the future breeding populations. In this study, weights were empirically determined from the nature of the data set and the breeding program requirements. Weights were kept constant throughout the analyses from AgMate and MateSel. The small progeny population required stringent penalizing weights on the coancestry (*w*_2_ = −10) and the additional constraint of not allowing any mate-pairs to have a coefficient of coancestry of more than 0.075. This helped the resulting optimal mating lists have a controlled increase in coancestry with a significant increase in genetic gain. One setback of determining the weights empirically is that it is somewhat arbitrary and depends on the breeder. To make the whole process of the candidate selections, determination of their contributions, and mating decisions more dynamic and tactical (Kinghorn & Shepherd, 1999), it is recommended that breeders run several analyses with varying combinations of weights to determine the suitable set of weights for the components that help meet the breeding criteria. Further research should include a more robust method of weight determination and optimization.

While both software, AgMate and MateSel, work efficiently to provide the optimized mating lists desired by the breeders under the same conditions, they do not produce the same optimum solution even though they utilize a similar differential evolution algorithm (DE). DE is an evolutionary algorithm that does not guarantee to provide the global optimum solution every run but is one of the most effective global optimization methods (Storn and Price 1997). No two runs under identical settings will result in identical results, i.e., running AgMate several times with the same inputs will not produce the same mating list for each run. The optimization process still provides optimum solutions, albeit it can be local or global optimums. Despite this caveat, the DE algorithm is robust in optimizations and will output the desired results every time. Numerous studies in the literature have shown and utilized the robustness of DE in optimizing objective functions (Carvalheiro et al., 2010; Kinghorn, 2011). This explains the difference in the results by AgMate and MateSel. The differences in the results between the two software can be accounted to different functionalities, like converge method, termination process, etc.

A licensed MateSel has advanced applications and more options than AgMate. There are many functionalities put together for animal breeding in MateSel. However, such advanced software with many options could be complicated for a non-computationally inclined breeder. Tree breeders need to be aware of such functions and be careful to avoid mistakes for a successful run on MateSel to create the desired optimized mating list. The manual for MateSel is 120 pages long and though not all the information is essential for monoecious tree breeding, understanding all the required inputs is a learning curve. This makes using MateSel, even with its front-end software, difficult to use effectively on the first try. While few of the functionalities are free, a licensed version is required to run the whole software, especially to take advantage of the monoecious functionality. MateSel is a front-end-only software, and changes are not allowed to the way the algorithm and analysis are run. Users can only make changes to the parameters provided as options. MateSel is also written in Fortran, a language that, even though very powerful, is not as popular in present times, making it hard to incorporate the next generation of tools and advanced algorithms.

AgMate is open-source software. AgMate is written in R and available on GitHub, allowing users the flexibility to edit the program to their necessity. Adding or removing functionalities is as easy as a click. For example, currently, the algorithm only optimizes based on EBVs, just a singular trait number, for genetic gain. Edits to the optimization equation or the input parameters can be made such that more traits of interest can be added to optimize based on all those parameters. R has been the prominent language in recent years, making it easy for users to understand the code. The availability of ShinyApp, a front-end for R software, also makes it easy to use the software from the front-end if a user is not well-versed in coding or even makes changes to the front-end allowing users to fit the software to their needs. With years and research, MateSel offers more functionalities for breeders than AgMate. While AgMate simplifies the creation of an optimized mating list and is user-friendly software, there is much room for improvement to help meet all the needs of the breeders.

Compared to traditional practices of selection based exclusively on genetic gain, this study uses advanced open-source software which incorporates state-of-the-art optimizing methods. This software showed the possibility of reducing the increase in coancestry to desired levels and minimizing inbreeding without compromising the expected genetic gain of the future progeny. The successful application of an advanced algorithm in two different software, which was originally created for different species, shows the flexibility of the algorithm. Future direction can incorporate more components to the objective function, such as minimum winter temperatures of selection, to stretch the flexibility of the software and have more breeding decisions handled by optimization algorithms.

AgMate and MateSel allowed a significant reduction in coancestry and inbreeding. AgMate performed better than MateSel in reducing coancestry (Figure3), whereas MateSel could more efficiently eliminate progeny inbreeding in most scenarios (Figure 4). This result can be associated with the frontier curve optimization finding method and balancing strategy functionalities adopted by MateSel, providing more opportunity for the software to find alternative candidates and mates to optimize the objective function and attain better outcomes for the components of the objective function. While MateSel was adopted from the field of physics and adopted for animal breeding and had many years of improvements under its belt, the comparable results from AgMate highlight the importance of adopting new algorithms and technologies from various unrelated fields to help in the advancement of breeding programs and their sustainability for the future. It is important to emphasize that the empirical commonly used strategies for controlling inbreeding in tree breeding, such as restricting the number of selected animals per family and not allowing full-sibs and half-sib mates, can be helpful at the beginning of the program but not effective in the long-term. The next advancement to incorporate into the optimizing AgMate software would be the use of genomic relationships derived from shared DNA markers, which have been shown to help better manage relatedness and inbreeding (Pryce, Hayes, and Goddard 2012).

**Figure 4.**
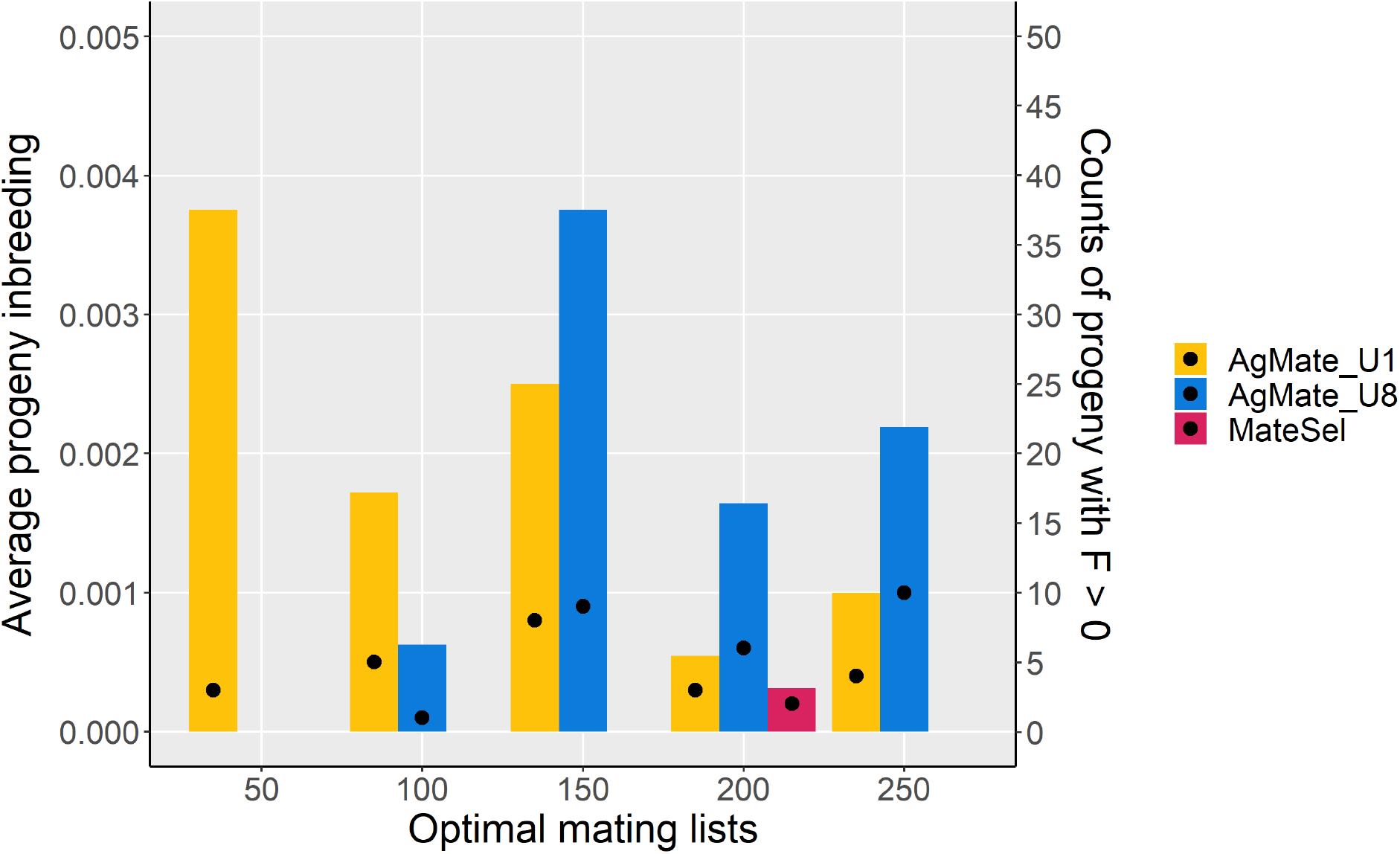
Comparison of AgMate and MateSel for progeny inbreeding levels under different mating lists (50 to 250). Three scenarios are shown: AgMate with MaxUse set at 1 (AgMate_U1) and 8 (AgMate_U8), and MateSel with MaxUse set at 8. Proposed progeny from AgMate showed average inbreeding levels, but these were limited to only a few individuals (1 to 10) contributing to the inbreeding in the progeny population. The number of progenies with inbreeding more than zero from the entire proposed mating list is shown by a black dot.

**Figure 5.**
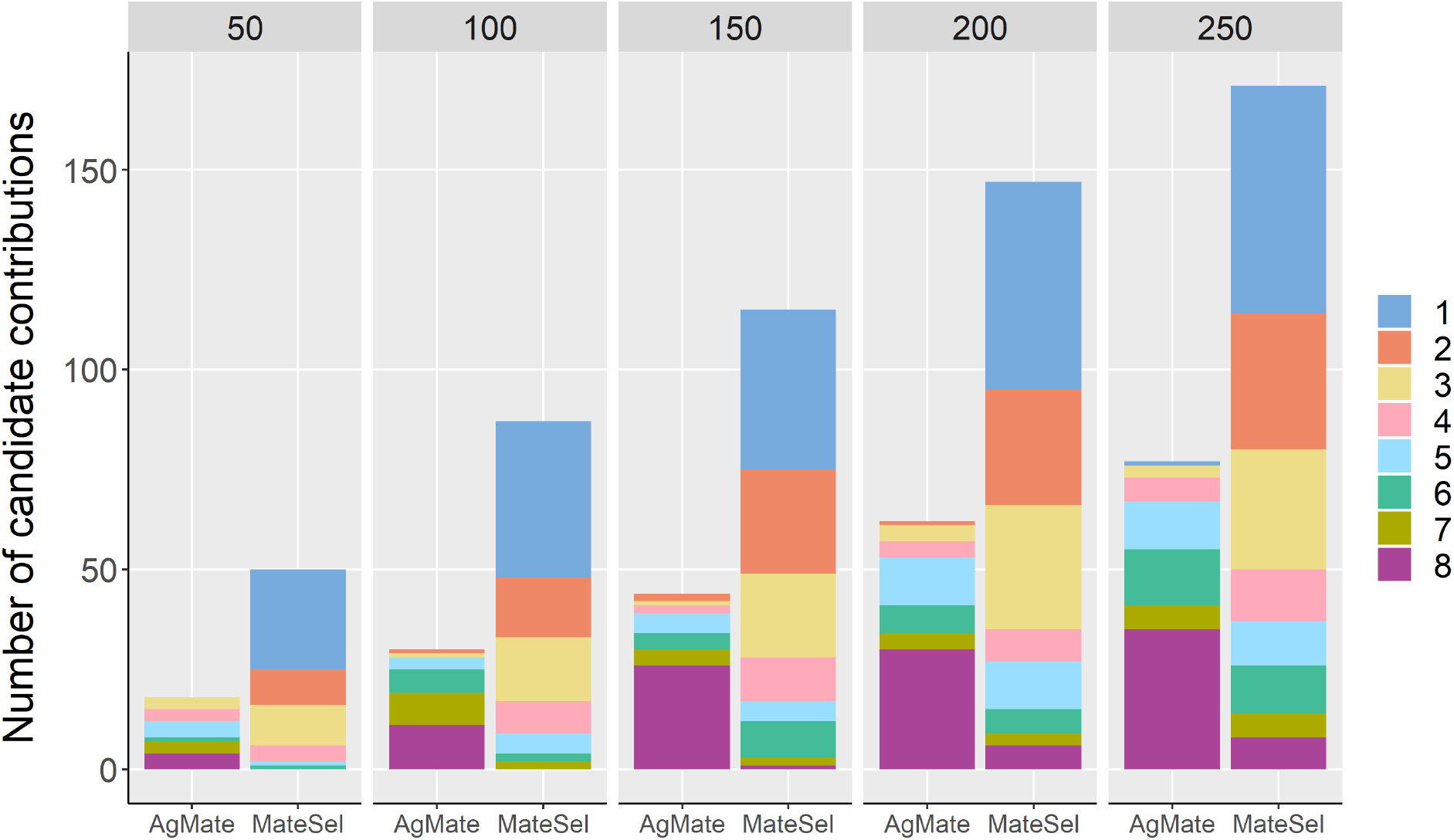
Candidate contributions in AgMate and MateSel analysis when MaxUse is set at 8 for different sizes of the mating list (50 to 250). AgMate uses fewer candidates than MateSel when MaxUse is greater than 1.

## Conclusion

The AgMate software was computationally efficient and flexible for practical applications in tree breeding. In contrast with empirical procedures for controlling inbreeding, the expected consequence of using the algorithm is to manage inbreeding more efficiently and promote higher genetic progress in the long term. Validating AgMate with a routinely used software like MateSel by Kinghorn showed the effectiveness of utilizing state-of-the-art algorithms like the differential evolution algorithm. AgMate is an easy-to-use and open-source software that simplifies difficult-to-comprehend algorithms. The usefulness and applications for software such as AgMate are profound for not just conifer breeding but also other monoecious species.

## Data archiving statement

The datasets, inputs, and outputs for AgMate and MateSel analyses can be found at https://doi.org/10.5061/dryad.tdz08kq2r. AgMate R code is available at https://github.com/khushigoda/AgMate. The ShinyApp version of AgMate is here: https://khushigoda.shinyapps.io/AgMate/.

## Acknowledgments

We would like to thank the staff and the members of the North Carolina State University Cooperative Tree Improvement Program for the *Pinus taeda* L. breeding population and their insights and recommendations for the AgMate software. We also thank Dr. Amanda Hulse-kemp, Dr. Christian Maltecca, and Dr. Gavin Conant for their feedback on the AgMate software. Goda would also like to thank Selene Schmittling and the data analysis consulting facility at NC State for their advice on resolving the issues in the AgMate software.

